# Visual detection of seizures in mice using supervised machine learning

**DOI:** 10.1101/2024.05.29.596520

**Authors:** Gautam Sabnis, Leinani Hession, J. Matthew Mahoney, Arie Mobley, Marina Santos, Vivek Kumar

## Abstract

Seizures are caused by abnormally synchronous brain activity that can result in changes in muscle tone, such as twitching, stiffness, limpness, or rhythmic jerking. These behavioral manifestations are clear on visual inspection and the most widely used seizure scoring systems in preclinical models, such as the Racine scale in rodents, use these behavioral patterns in semiquantitative seizure intensity scores. However, visual inspection is time-consuming, low-throughput, and partially subjective, and there is a need for rigorously quantitative approaches that are scalable. In this study, we used supervised machine learning approaches to develop automated classifiers to predict seizure severity directly from noninvasive video data. Using the PTZ-induced seizure model in mice, we trained video-only classifiers to predict ictal events, combined these events to predict an univariate seizure intensity for a recording session, as well as time-varying seizure intensity scores. Our results show, for the first time, that seizure events and overall intensity can be rigorously quantified directly from overhead video of mice in a standard open field using supervised approaches. These results enable high-throughput, noninvasive, and standardized seizure scoring for downstream applications such as neurogenetics and therapeutic discovery.

## 2 Introduction

Epilepsy is a broad collection of neurological disorders defined by the occurrence of spontaneous, unprovoked seizures, which are often comorbid with other neurological disorders, including autism spectrum disorder, anxiety, and intellectual disability [1, 2]. Seizures are caused by abnormally synchronous brain activity that can result in changes in muscle tone, such as twitching, stiffness, limpness, or rhythmic jerking, and loss of consciousness [1]. Genetics has a substantial influence on epilepsy, including both common variants that modify risk for common epilepsies [3] and monogenic etiologies that cause severe developmental and epileptic encephalopathies [4, 5, 6, 7, 8]. With these findings, the potential for genetically informed, precision therapy is beginning to be realized [9, 10]. However, despite high heritability and the many known monogenic genetic etiologies, there is large unmet medical need in the treatment of patients; nearly one third of patients are refractory to all existing medications and there are no known disease-modifying therapies [11]. Thus, mouse models of epilepsy are a critical experimental resource, given our ability to control mouse genetics and the high homology between mouse and human neural circuits underlying seizures. However, maximal use of mice for quantitative genetics and pre-clinical modeling requires rigorous, quantitative, high-throughput seizure phenotyping.

Clinically, seizures are diagnosed using an electroencephalogram (EEG), which measures the electrical activity in the brain and can detect abnormal synchronous activity as large rhythmic patterns in brain activity. However, the use of EEG in pre-clinical studies has significant limitations. In particular, EEG is invasive, expensive, and low throughput, requiring surgery to implant a monitoring device. However, convulsive seizures in animal models have behavioral manifestations that correlate with underlying brain activity, and therefore with seizure severity, although these correlates can depend on species and etiology [12, 13, 14]. Thus, there are several options for scoring seizure severity that differ in their precision and quantitative rigor. The simplest approach is to observe seizure events, either in real time or on video, and score them according to a standard rubric, such as the Racine scale [12, 13]. This is the lowest cost option but is not high-throughput enough for long-term monitoring and is partially subjective. In contrast, invasive EEG monitoring yields direct measurements of brain activity and has been the gold-standard for seizure studies, but these systems always require invasive long-term implantation of electrodes into or near the brain, which carries a non-trivial risk of harm to the animal via infection and can alter motor behavior and circadian rhythms [15, 16, 17]. Furthermore, depending on the recording system (e.g., tethered vs. wireless telemetry), the equipment itself may limit the other types of behavioral screening that can be performed in those animals. This can be a significant limitation for studies that seek to correlate seizure activity with other behavioral changes to study, for example, the well-established relationship between epilepsy and autism [1]. Low-profile, wireless systems impose the fewest restrictions on other behavioral assays (see Lundt *et al.* and references therein [15]), but are also the most expensive, which also limits throughput for most investigators. An intermediate alternative is to use a minimally invasive or non-invasive surrogate signals, such as accelerometry [18], to allow much higher throughput and lower costs, along with rigorous signals for reproducible seizure detection. In all the above approaches, seizures are scored by an expert analyst, perhaps after a selection of putative seizure video clips by auxiliary signal processing. However, to date, there have been no approaches to directly score seizure intensity in animal models from video data directly.

Recent advancements in animal behavior annotation have resulted from applications of new methods from the statistical learning and computer science fields for biological tasks. These combine machine learning and computer vision (machine vision) and are often referred to as "artificial intelligence (AI)" [19, 20, 21, 22]. We have successfully applied these methods for tracking [23], action detection [24], frailty [25], nociception [26, 27], sleep [28], and gait and posture analysis [29]. Furthermore, we have built a shared open platform for end-to-end behavior annotation [30]. Given these advances in machine vision, we explored whether a video-only computer vision approach could be a solution for non-invasive seizure scoring in mice.

Gschwind *et al.* have recently used an unsupervised approach to detect behavior anomalies due to seizures [31]. Their method, MoSeq [32], automatically classifies video into discrete sequences of "syllables", each of which is a short behavioral pattern (*∼* 300 milliseconds). Interestingly, while patterns in the MoSeq syllables did correlate with the severity of seizure events, the classification performance was modest (*F*_1_ = 0.39 *±* 0.13), which they attributed to several factors, including that MoSeq syllables correspond to very short behaviors lasting only a few hundred milliseconds, while human scoring is based on highly specific behaviors lasting much longer [31]. Thus, the most robust changes observed using unsupervised methods are interictal rather than ictal events. These findings, combined with our prior work utilizing supervised and heuristic classification of rodent behaviors, such as grooming [24], and combining multiple behaviors to score higher-level constructs such as frailty [25] and nociception [27] strongly suggest that the behavioral manifestations of frank seizures should be robustly detectable from video data, but no studies have addressed automatically scoring seizure severity *per se* in animal models. This is a significant gap because long-term monitoring of large cohorts may be critical for future pre-clinical studies of epilepsy.

In this study, we used a supervised learning approach to detect seizures in mouse behavioral videos. The community standard seizure severity score for pre-clinical models is the Racine scale[12, 13, 14], which is an ordinal scale from one to seven denoting progressively more pronounced behavioral manifestations of seizures, from whisker trembling (Racine score = 1) to a tonic-clonic seizure followed by tonic extension and possibly respiratory arrest or death (Racine score = 7) [13]. In order to enrich for seizure events we induced seizures using the convulsant pentylenetetrazole (PTZ), a gamma-aminobutyric acid (GABAA) receptor antagonist, has been commonly reported in the literature [33, 13, 34]. We hypothesized that we could train robust, automated classifiers to predict seizure severity using video data alone. Overall, we show that supervised methods can accurately detect specific seizure events and that these events can be combined to create a seizure intensity score.

## 3 Results

The design of our experiment is illustrated in Figure 1A. To create robust training data with many seizures, we challenged our mice using PTZ at varying doses. We monitored mice in an open field during PTZ-induced seizures that were scored according to the Racine scale by an expert observer [13]. This created our human-annotated training data for seizure intensity. In order to automate the seizure detection, we used the following steps. First, we extracted pose-estimation and body-segmentation time series from raw behavioral videos using existing deep neural network models [30, 29, 23]. Second, from the extracted time series, we built classifiers for six characteristic behavioral seizure manifestations: freezing (*i.e.*, behavioral arrest, score of 1), Straub tail (score of 4), leg splaying (score of 4), circling (score of 5), side seizure and wild jumping (score of 6). Third, and finally, we summarized the classifier outputs as a set of features (*e.g.*, "time spent in side seizure") that were used in an ordinal regression model to predict seizure severity. To detect seizure intensity, we carried out two analyses. In a first analysis, we predicted the seizure severity over the course of an entire session. In a second analysis, we predicted time-varying seizure intensity.

**Figure 1:**
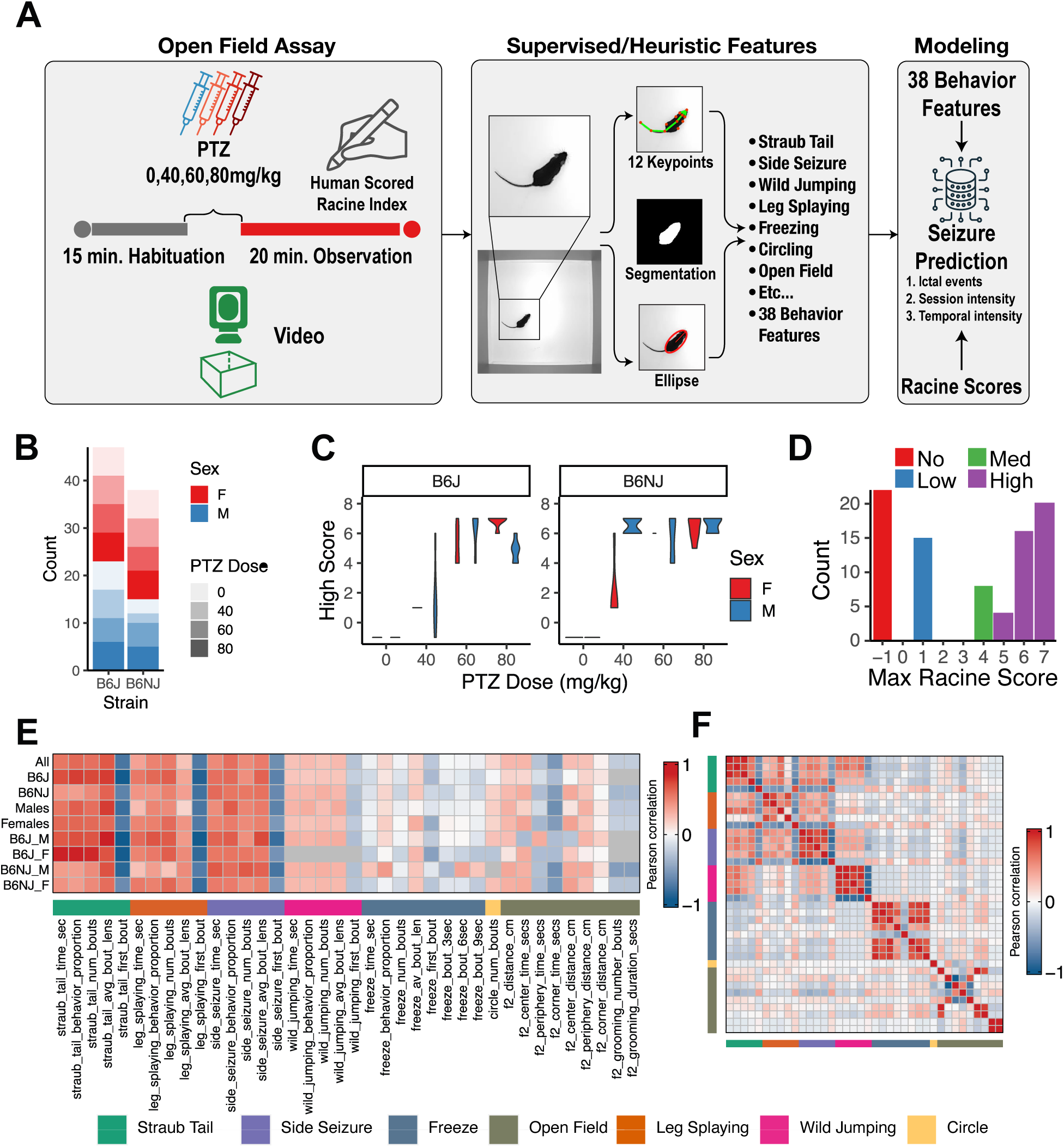
Experimental design, manual Racine scores, and behavioral features for seizure classification. (A) Visual representation of the experimental workflow (B) A bar plot summarizing the number of animals per strain, sex, and PTZ dose. (C) A violin plot summarizing the human-annotated data. Racine scores range from -1 to 7. (D) A histogram distribution of the scores (ground truth) assigned by the tester. The colors indicate the Racine intensity group from Table 1 (E) Pearson correlations of features’ summary statistics with Seizure (see Table 1). (F) A correlation matrix showing the estimated bi-variate correlations across features.

**Table 1:** Racine Scale scores and behaviors.

| Score | Description | Seizure Intensity |
| --- | --- | --- |
| -1 | Normal baseline | No |
| 0 | Whisker trembling | Low |
| 1 | Sudden behavioral arrest | Low |
| 2 | Facial jerking | Low |
| 3 | Neck jerks | Medium |
| 4 | Clonic seizure (sitting) | Medium |
| 5 | Clonic, tonic-clonic seizure (lying on belly) | High |
| 6 | Clonic, tonic-clonic seizure (lying on side) and wild jumping | High |
| 7 | Tonic extension, possibly leading to respiratory arrest and death (laying down with forelimbs and hindlimbs outstretched) | High |

### 3.1 Manual seizure scoring

We used both male and female C57BL/6J (B6J) and C57BL/6NJ (B6NJ) mice with 4 doses of PTZ – 0 mg/kg (vehicle), 40 mg/kg, 60 mg/kg, and 80 mg/kg (Figure 1B). These doses, strains, and methods were based on a previously published paper [13]. Mice were allowed to acclimate for at least 15 minutes in an open field arena before being injected with PTZ and returned to the open field (Figure 1A). Mice were observed by a single trained observer for a maximum of 20 minutes and given a Racine score. JAX IACUC protocols dictate that any mouse observed to have tonic extension or maintained a Racine score greater than 3 (neck jerks) for one minute must be immediately euthanized. A mouse is also removed from the open field and euthanized following the first signs of tonic extension or extreme seizure activity. Following video capture, data were processed using machine learning and computer vision methods described below.

Racine scale scores (1 through 7) were assigned to behaviors during the session, with a score of -1 assigned to baseline, non-seizure activity by the observer. When mice were presented with multiple Racine scores, the highest score was used as an overall score (all Racine scores are in Table S1). The distribution of Racine scores across strains and aggregate distribution showed that across both strains of mice, the Racine score increased as a function of dose (Figure 1C, D, S1). We binned the Racine scores into 4 categories of seizure intensity for downstream modeling (no, low, medium, and high, Table 1 and Figure 1D).We observed all modalities of seizures (Table S1) and after taking the maximum Racine score for each animal, we expected that it would be difficult to distinguish the medium group from the rest due to the uneven distribution of Racine scores (Figure 1D) and thus seizure levels. This manual Racine scoring analysis indicated that the data were of high quality and suitable for modeling ictal events.

### 3.2 Seizure prediction

To predict seizure events, we developed classifiers for six seizure-associated behaviors: Straub tail, leg splaying, side seizure, wild jumping, freezing, and circling (described in Table 2, [35, 36, 37, 13]). Examples of these behaviors can be seen in Supplementary Videos 1-4. These behaviors were chosen based on whether the behavior was easy to distinguish on video and whether there were enough clear instances of that behavior to train a classifier. The JABS active learning module uses pose-based features extracted from the videos to classify behaviors [30, 38]. We used JABS to classify Straub tail, leg splaying, side seizure, and wild jumping events. To classify freezing and circling, we define two heuristic classifiers (Table 2). Overall, all seizure associated behavior classifiers had good performance based on *F_β_* (Table 2). Next, we extracted summary statistics for each behavior, including total behavior time, proportion of behavior time in trial, number of behavioral bouts, and average length of behavioral bouts similar to our previous applications ([27], fully described in Table S2). Combined with previously used open field features, we used 38 total features.

**Table 2:** Ictal behaviors descriptions and classifier accuracy. Supplementary videos are also indicated.

| Type | Behavior | Description | $F_\beta$ |
| --- | --- | --- | --- |
| JABS classifier | Straub tail | Tail stiffens, rises quickly, may jerk down (Supp. Video 1) | 0.85 |
| JABS classifier | Leg splaying | Hind legs straighten, visible, wide stance (Supp. Video 2) | 0.94 |
| JABS classifier | Side seizure | Mouse falls to side, shakes/seizes (Supp. Video 3) | 0.79 |
| JABS classifier | Wild jumping | Mouse runs and jumps quickly around arena (Supp. Video 4) | 0.79 |
| Heuristic | Freezing | No movement for at least 3 seconds |  |
| Heuristic | Circling | Tight circular locomotion, followed by another within 6 seconds |  |

Next we determined if these features are useful for seizure intensity classification. As a preliminary test for association to seizure intensity, we correlated these summary statistics with seizure groups and PTZ dose for all test groups, including sex and strain stratification (all mice, all females, all males, all B6J, all B6J females, all B6J males, all B6NJ, all B6NJ females, and all B6NJ males (Figure 1E and Supplementary Figure S1A). We found certain behaviors strongly correlate with PTZ dose and seizure intensity groups. For example, the amount of Straub tail behavior had consistently high positive correlation with dose across all strains and sexes. Straub tail and side seizure behavior also showed positive correlations with dose, but with less significance for B6NJ. Certain seizure behaviors such as wild jumping and circling were rare in our dataset and did not show a strong correlation with dose or seizure intensity (Figure 1E, pink and yellow), mainly because there were very few instances of these in our data. Freezing behavior is quite frequent, however it occurs in all three PTZ doses, and then does not show a strong correlation with dose or seizure intensity (Figure 1E, navy blue). We also calculated the proportion of time across all total behaviors relative to the time in the arena for each mouse, which showed a significant positive correlation with the dose for all groups except B6NJ males (Figure 1E). Thus, the correlation heatmap for behavior summary statistics with highest Racine score is very similar to the one for PTZ dose (Supplementary Figure S1A). In contrast to seizure specific behaviors, routine open field behaviors, such as activity and anxiety measures, do not correlate with the intensity of the seizures or the dose of PTZ (Figure 1E, dark green).

The correlation matrix among all features shows multi-collinearity, with extremely strong correlations among features derived from a single behavior (*e.g.*, leg splaying) and substantial correlation between different behaviors (Figure 1F). Thus, these behaviors co-vary across animals in structured ways, as expected for highly stereotyped seizures. This can cause model parameters estimated from the data to be unreliable and, in some cases, even unidentifiable. To circumvent this, we use a regularized version of the model [39, 40].

### 3.3 Visual detection of seizures

We used the seizure behavior classifiers to visualize individual seizure events over time with ethograms (Figure 2). We plotted all seizure behaviors for each mouse at each dose (Figure 2A), and for each behavior at each dose (Figure 2B). Since mice were removed if they had severe seizures or a period of mild seizures, we mark the end of each animal’s test (black points). We observe that as the dose increased, mice were removed from the arena in less time, which indicates earlier onset of severe seizures. Most of the animals in the 80 mg/kg dose were in the arena less than 4 minutes before being euthanized due to severe seizures. We also observe that mild events such as freezing and Straub tail are observed in the 40mg/kg dose. At 60 and 80mg/kg we observe all classes of seizures, including wild jumping, side seizures, circling, and leg splaying. Overall, these results visually show the onset, frequency, and euthanasia of each animal over time. It also demonstrates that mild and severe seizure events are captured by distinct behaviors – freezing and Straub tail behaviors, vs. leg splaying, circling, side seizures, and wild jumping, respectively. This validates our approach of scoring individual seizure events using a supervised approach.

**Figure 2:**
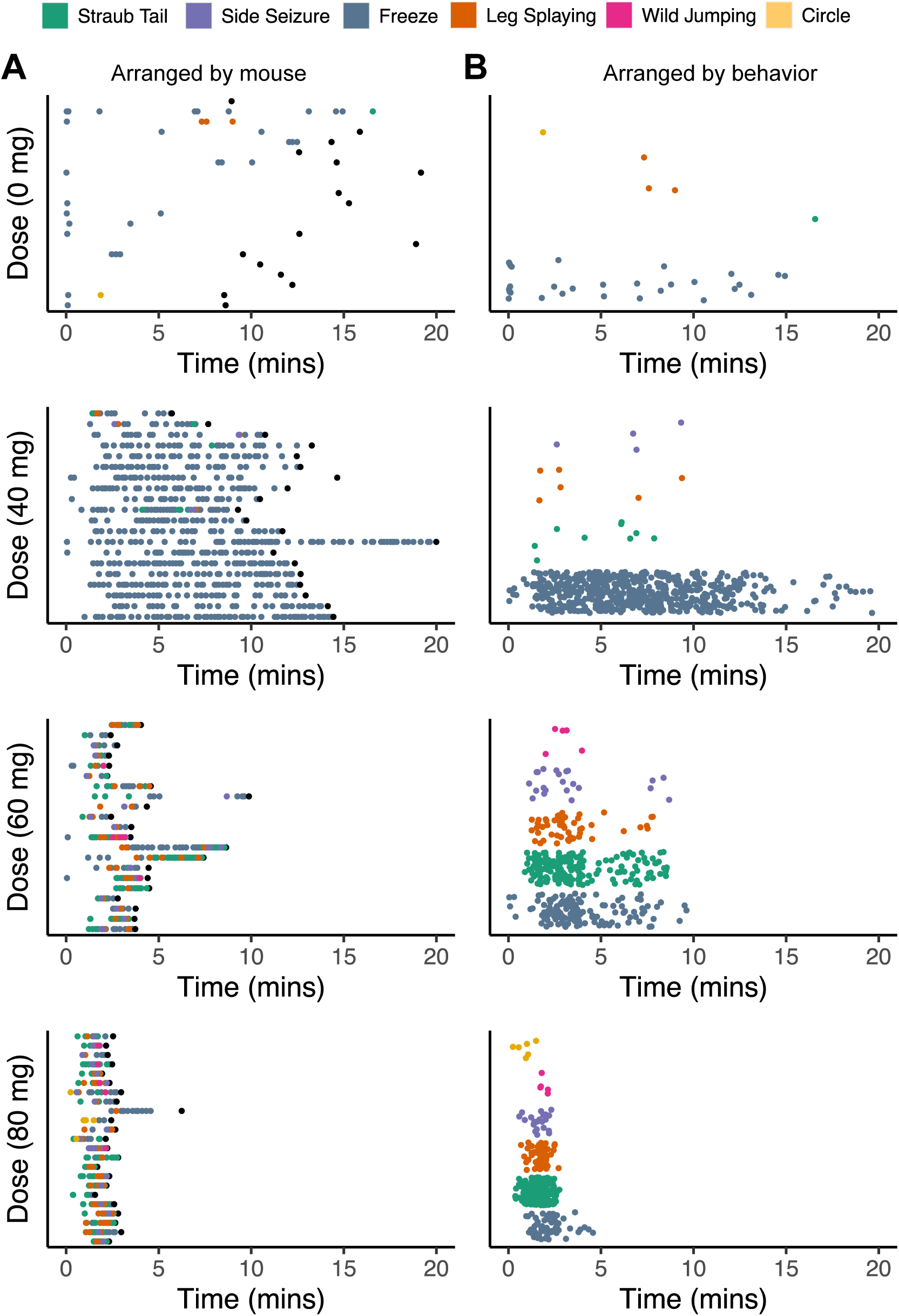
Ethograms of seizures behaviors. Time 0 represents time when mice are placed in open field arena after injection with vehicle (0mg) or increasing dose of PTZ. (A) Each dot represents the occurrence of a behavior (all colors except black) for a mouse (each row on y-axis) at a certain time point (x-axis) during the duration of the experiment before the mouse is removed (black dots). (B) Same as (A) except all behaviors performed by every animal are arranged by behavior rather than animal.

### 3.4 An interpretable univariate seizure scale

Next, we sought to create a univariate scale using the 38 features. Univariate scales can be a powerful method to condense multiple features into one metric that can be applied towards multiple tasks. They are frequently used in biological applications [41] and can be constructed using various regression modeling approaches. A univariate seizure scale offers several advantages: 1) easy to construct once the model is fit; 2) it is highly interpretable; and 3) it can easily be implemented with new data. We have applied it previously for frailty indexing and pain scoring [25, 27].

We first established that our features capture differences that distinguish animals given different doses and belonging to different seizure groups based on their Racine scores (Figure 3A,B). Linear discriminant analysis (LDA) indicated that both dose and seizure intensity could be distinguished. We find that the medium and high intensity Racine groups and the 60 and 80mg/kg PTZ dose groups are challenging to distinguish (Figure 3A,B, green and purple). This indicates that even the 60mg/kg dose is likely saturating our dose response curve.

**Figure 3:**
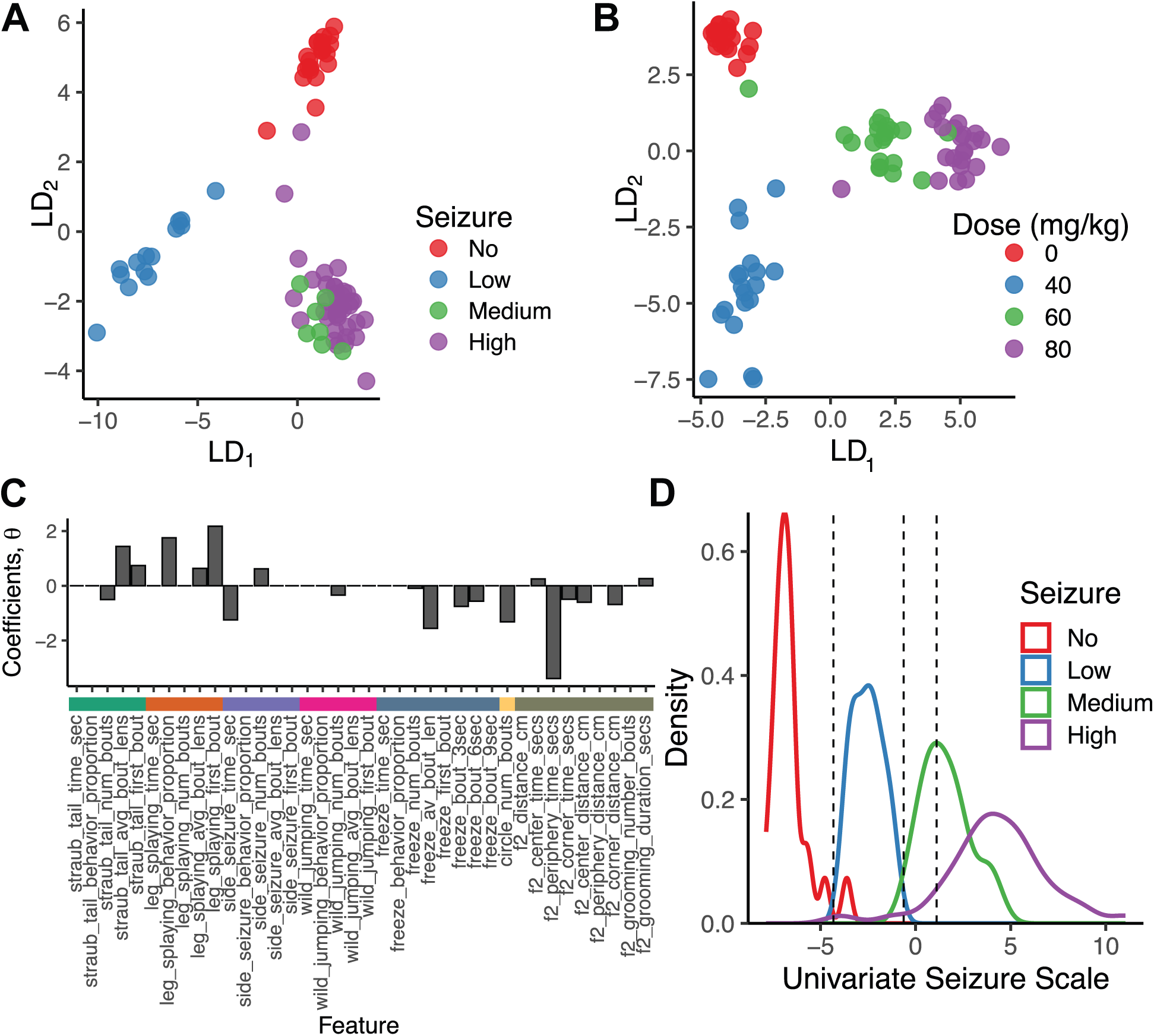
Univariate seizure scale for seizure intensity (A) We used linear discriminant analysis (LDA) to quantitatively distinguish across animals (A) belonging to different Racine groups, (B) administered different dosages. (C) We have plotted the features’ contributions to the seizure intensity scale. (D) We created a univariate seizure scale by training ordinal regression models to create a latent continuous scale with segments corresponding to different seizure intensities. The resulting distribution of mice along the scale shows an ideal separation of seizure intensity in each segment.

To construct a univariate scale, we first trained a supervised latent variable model that utilizes features to predict seizure intensity. Specifically, we used an ordinal regression model [42] to estimate the probability of each subject being classified as having no, low, medium, or high seizures. We adopted the elastic net penalty [40] for regularization to address multicollinearity (Figure 1F), perform feature selection, and improve prediction accuracy. We used 10-fold nested cross-validation to tune the regularization hyperparameter and get an unbiased estimate of the tuned model’s out-of-sample misclassification rate. Averaging over the folds gave us a misclassification error of 0.17*±*0.13. For comparison, a classifier that guesses seizure intensity would have a misclassification error of 0.75. We have plotted the estimated feature weights, where a positive (or a negative) coefficient shifts probability towards a higher (or a lower) seizure group (Figure 3C). Indeed, we see positive coefficients estimated for both Straub tail and leg splaying (number of bouts and average bout length); even side seizure has an estimated positive coefficient. We found most features associated with freezing to have negative coefficients, with only more prolonged bouts of freezing (9 seconds) having a positive coefficient. We used the feature weights estimated from the model to construct a univariate seizure scale by taking a weighted sum of the features using the estimated model parameters as weights. The scale allowed us to distinguish animals with no and low seizures clearly from medium and high seizures, although distinguishing animals with medium levels of seizures from high seizures is more challenging (Figure 3D). As a baseline comparison, we trained a supervised latent variable that utilizes features to predict the Racine score on the original scale. We found the average misclassification error rate using 10-fold cross-validation to be 0.43 *±* 0.14 (Figure S1C,D). We reason this is likely due to human subjectivity in Racine scoring.

### 3.5 Temporal detection of seizure intensity

Our results above quantify seizure severity by considering a complete seizure event. However, seizures are caused by a sudden and uncontrolled burst of synchronous electrical activity in the brain and generally occur spontaneously, evolve in time, and, in many rodent models, can be rare events [1, 43, 44]. Thus, beyond scoring complete events, which can last several minutes, it is desirable to detect the temporal variation in seizure intensity as a function of time, both to identify seizure events in potentially long recordings and to track the seizure evolution. Thus, we sought to build a temporally resolved seizure intensity scale that operated on a minute of data that could potentially be used for temporal seizure detection (Figure 4A). Using the PTZ dataset from above, we trained an ordinal linear mixed model [45] to predict the overall seizure intensity based on 1-minute bins of data from each mouse. Our underlying assumption was that characteristic 1-minute bins would correlate with the evolution of seizure intensity.

**Figure 4:**
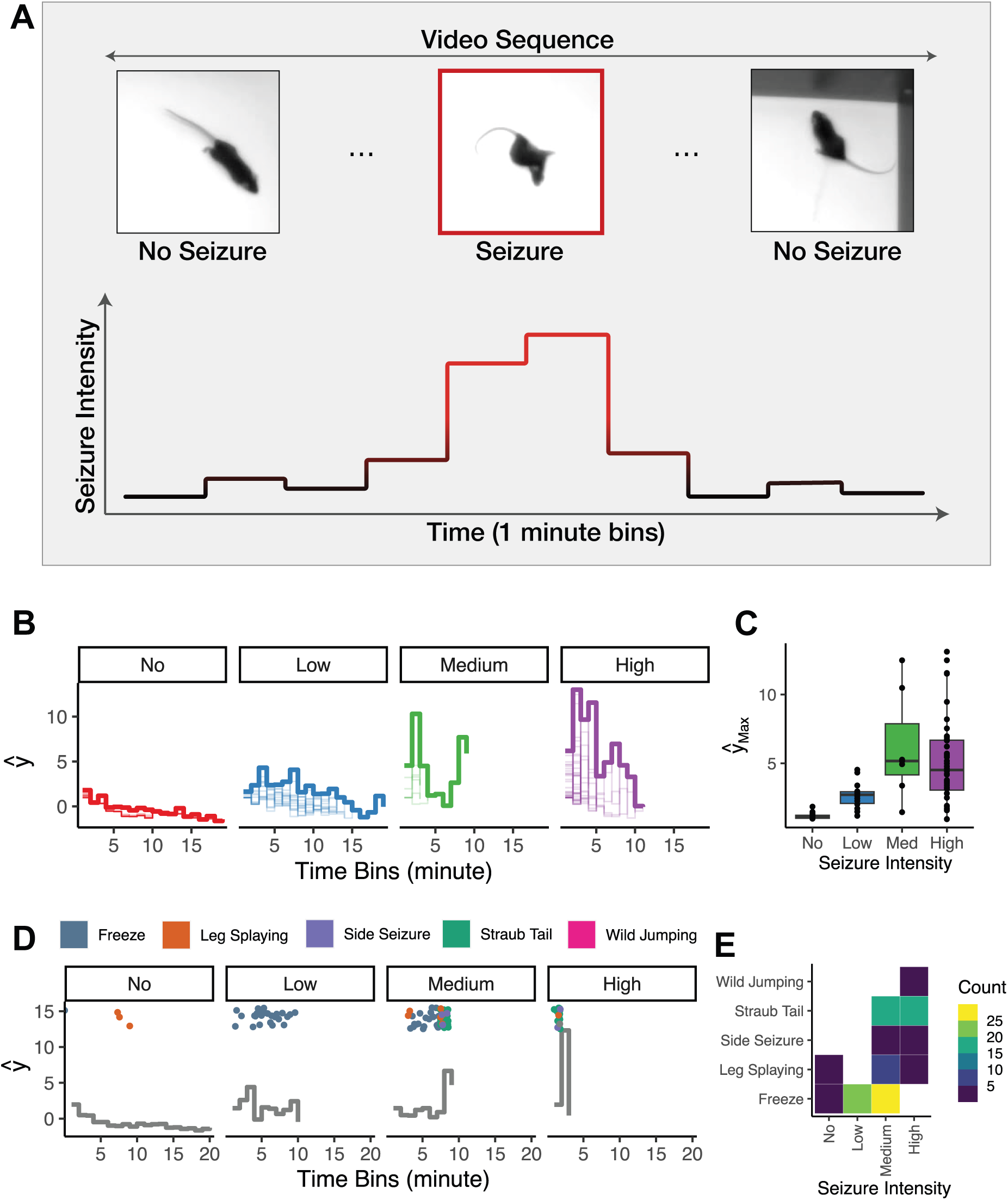
Temporal analysis of seizure intensity. (A) Seizure intensity score is assigned on one-minute video segments. An ideal seizure detector will identify rare time bins when seizures occur (e.g. red frame grab). We use a weighted sum score of features at each 1-minute interval for this analysis. (B) Instantaneous seizure intensity estimates across 20 1-minute intervals were estimated using supervised hierarchical regression models. The thin lines correspond to individual mice seizure profiles while the bold lines are the maximum seizure intensity for each seizure group. Also see Figure S1. (C) Validation of the instantaneous seizure intensity model using leave-one-animal cross-validation (LOOCV) where we estimated the maximum seizure intensity of held-out mice (y-axis) and compared it to their true Racine groups (x-axis). (D) We select representative animals from each group and overlay annotated behaviors on top of predicted behavior profiles. (E) Temporal analysis correlates with intense seizure events. We count the number of bouts (color) of annotated behaviors (y-axis) in each seizure group (x-axis) for the animals in D.

We undertook quantitative and qualitative approaches to validate our model on previously unseen data. We removed one animal to perform leave-one-out cross-validation (LOOCV) and validated the model’s performance on the unseen animal. In particular, we regressed the seizure intensity on the features by treating the repeatedly measured features across each minute as a random effect. We estimated the model parameters from the data. We then computed the predicted seizure intensity score for the left-out animal as a function of time, attempting to approximate a behaviorist labeling the videos and assigning a Racine score at each 1-minute bin (Figures 4B, S1E). We validated the model by calculating the predicted temporal profile’s mean, median, and maximum measures. As with the univariate scale, we found that the three summary measures captured group differences between the seizure intensity groups (Figure 4C, Supplementary Figure S1F). In other words, the average temporal profiles recapitulated the overall group differences. As before, the predicted temporal profiles did not distinguish as strongly between medium and high seizure groups as the other groups.

The maximum measure identifies the minute bin at which we predicted each left-out animal to have the maximum score. We picked one animal from each seizure intensity group with the highest predicted score and overlayed the behavior classifier predictions (Figure 4D). We found that the predicted temporal profiles coincided with the co-occurrence of seizure-specific behavioral bouts (Figure 4E), demonstrating a qualitative agreement between the seizure intensity profiles and major seizure events.

## 4 Discussion

In this study, we used supervised machine learning to develop a new model for automated seizure scoring using open-field video monitoring. Specifically, we perform three levels of analysis. We are able to detect specific seizure behaviors. We use these and other features to predict the session-level maximum Racine score. Finally, we carry out time-resolved seizure intensity scoring to detect seizure intensity in a time series data. Our approach is non-invasive, reproducible, and scalable.

The virtues of purely video-based monitoring are appreciated by the epilepsy field as an alternative to cumbersome gold standard EEG recordings. Moreover, outside of epilepsy, it is appreciated that seizures are a significant class of adverse events leading to high attrition in drug development outside of seizure disorders, and automated video-based seizure scoring is recognized as a potential solution [46, 47]. While this approach has been applied towards detection of seizures in humans [48, 49, 50, 51, 52, 53, 54, 55], and is recognized as a critical frontier for human phenotyping [56], visual detection of seizures in rodents has not been extensively explored. Gschwind *et al.* recently reported that unsupervised segmentation of sub-second behavioral motifs was sufficient to discriminate between epileptic and non-epileptic animals [31]. Their approach utilizing MoSeq [32] uses inter-ictal behaviors to carry out this task. While MoSeq was able to detect behavioral differences between epileptic and non-epileptic animals, the detected behavioral motifs did not neatly classify into seizure and non-seizure motifs. In contrast, by using a supervised learning approach, we were able to directly classify a robust set of behaviors associated with seizures (Table 2) and integrate them into seizure intensity scores.

It is important to note that our approach is not a surrogate for Racine scoring *per se*, in that we do not classify all Racine score features. In particular, some of the semiological features in the Racine scale, such as facial twitches and whisker trembling, are extremely subtle and are not easily detected by overhead video. We focus on those features that involve relatively large changes detectable at the level of posture or locomotion. This approach dramatically simplifies the video recording set up to be in line with those for widely adopted behavioral tasks and, in particular, does not require special equipment such as a depth camera. Furthermore, a computational ethology [57] approach has the potential to enable detection of multiple behaviors simultaneously. Future developments can combine seizure detection with other behaviors such as homeostatic behaviors, social behaviors, and more to holistically phenotype an animal. Furthermore, unsupervised and supervised methods could be used in a complementary assay to provide rich partitioning of ictal and intericatal behaviors. New version of MoSeq enables behavior segmentation using keypoint data [58]. Other unsupervised methods such as MotionMapper, VAME, B-SOiD and others could be explored [59, 60, 61].

Our predictive models were particularly strong at separating out the "No" and "Low" seizure groups from each other and from the "Medium" and "High" groups, while it was harder to distinguish the "Medium" and "High" groups from each other. We attribute this largely to the performance of the base classifiers (Table 2), where the F1 scores for both "Side seizure" and "Wild jumping", which discriminate between the "Medium" and "High" groups on the Racine scale, were lower than the F1 scores for "Straub tail" and "Leg splaying". Thus, noise from the base classifier outputs potentially contributed to the overlap between the "Medium" and "High" groups in the feature space. However, we also note that the number of "Medium" scores was substantially smaller than the other three groups, and was composed exclusively of individuals with overall Racine scores of 4. Thus, discrimination of "Medium" from "High", in this case, required detecting the difference between a sitting tonic-clonic seizure (Racine score = 4) and a tonic-clonic seizure on the belly (Racine score = 5) (Table 1). Thus, increasing the number of individuals in the "Medium" class may improve future models, although we stress that titrating the dosage to achieve "Medium" and not "High" seizure responses is a significant experimental challenge. More generally, we argue that seizure severity is truly a continuum and that, while binning this continuum into discrete categories can aid in model training, we should always expect edge cases that are difficult to classify. Nevertheless, our model’s performance significantly captures the overall variability in seizure severity.

Taken together, our results demonstrate that the major semiological features of mouse behavioral seizures are robustly detectable from video data alone. Our models provide reproducible quantitative scores that we expect to be valuable for downstream applications, such as drug screening studies, quantitative genetics, and correlating seizure phenotypes with behavioral comorbidities.

## 5 Methods

### 5.1 Animals and PTZ protocol

In this study, we tested both male and female C57BL/6J (B6J, JAX 000664) and C57BL/6NJ (B6NJ, JAX 005304) mice with 4 doses of PTZ (pentylenetetrazole, CAS 54-95-5, Sigma, cat# P6500; 0 mg/kg (vehicle), 40 mg/kg, 60 mg/kg, and 80 mg/kg) injected intraperitoneally. We collected a total of 85 videos (47 B6J, 38 B6NJ) (Figure 1B). All doses had at least 5 males and 5 females, with the exception of B6NJ at 0 mg/kg (Figure 1B). 8-12 week old mice were shipped from JAX production to Kumar Lab testing rooms. They were allowed to habituate for at least 1 week prior to testing. Mice were weighed prior to testing. Fresh PTZ solutions were formulated in 0. 9% saline at appropriate doses and administered by intraperitoneal injections.

### 5.2 Open field assay

The open field behavioral assays were conducted as previously described using a top-down camera view [23, 62] using our JAX Animal Behavior System (JABS) [30]. Mice were allowed to habituate for at least 15 minutes in the open field arena before being injected with one of the listed doses of PTZ and returned to the open field. Following our IACUC-approved protocol, mice were directly observed by a tester for a maximum of 20 minutes and given a Racine score [13]. A mouse was removed from the open field and euthanized following the first signs of tonic extension or extreme seizure activity. A mouse was also euthanized if it maintained a Racine score greater than 3 (neck jerks) for a minute. Additionally, in the control group, if a mouse behaved normally for 5 minutes (i.e. no behaviors associated with seizures were observed), the animal was removed from the arena to save time from manual observation. Using this protocol, we collected a total of 85 videos (47 B6, 38 B6NJ) (Figure 1B). All doses had at least 5 males and 5 females, with the exception of B6NJ at 0 mg/kg dose (Figure 1B). A direct observer of the animals did the scoring.

### 5.3 Video recording, segmentation, and tracking

The JABS open field arena, video capture methods, and tracking and segmentation networks are as detailed previously [23, 30]. We use a neural network trained to produce a segmentation mask of the mouse to produce an ellipse fit of the mouse at each frame as well as a mouse track. JABS is a versatile open source system, which we have used for gait and posture measures [29], grooming [24], biological age [25], and pain states [27].

### 5.4 JABS classifiers

Briefly, our data acquisition uses custom-designed standardized data acquisition hardware and software that provides a controlled environment, optimized video storage, and live monitoring capabilities. This is leveraged by the JABS annotation and classification software which is a python based GUI utility for behavior annotation and training classifiers using the annotated data. Full details can be found in Beane et al. (2022) [30]. One can then use the trained classifiers to predict whether behavior happens or not in the unlabeled frames. For this study, we trained behavioral classifiers for behaviors such as straub tail, side seizure, leg splaying, and wild jumping (Table 2). Based on previous previous experience, we use bout wise accuracy metrics with an IoU of 50% [30].

### 5.5 Open field measures and feature engineering

Open field measures were derived from ellipse tracking of mice as described previously [23]. Tracking was used to produce locomotor activity and anxiety features. Freezing behavior was heuristically derived by taking the average speed of the nose, base of head, and base of tail points at each frame, and finding periods of at least 3 seconds where the average speed of the mouse was less than 0.01 cm/sec. For detecting tight circling events, we begin by observing changes in angles the mouse is facing. Once these changes in direction exceed 360 degrees in either direction, we segment out that as a circle event. We then calculate the distance traveled during that event. If the mouse has traveled more than 6 cm, then the mouse was walking around the arena. Distances less than 6cm indicate a tight circling event. To adjust for the different durations that animals were observed during experiments with varying PTZ doses, we standardized each feature by the time (in minutes) the animal was taken out.

### 5.6 Linear discriminant analysis (LDA)

We centered the features to have zero mean. To address multicollinearity across features and avoid overfitting with LDA [63], we performed principal component analysis (PCA) to obtain projections of features that are mutually uncorrelated and ordered by variance. We used all principal components (PCs) in the subsequent LDA algorithm to avoid the risk of throwing away critical dimensions. We projected the data to 2D LDA space for visualization.

### 5.7 Ordinal linear models for predicting seizure intensity

We used a supervised latent variable model (ordinal regression) to regress a continuous latent variable underlying the seizure groups (highest Racine scores) (see Table S1) onto behavioral features. This produced a univariate seizure scale divided into discontinuous segments based on the threshold parameters estimated in a data-driven manner by the model. More specifically, we used cumulative link models, a type of ordinal regression, most commonly used to analyze ordinal data [64]. The mouse’s seizure group, one of the four ordinal categories (High, Medium, Low and No) is regressed onto the mouse’s measured behavioral features. The regression coefficients and thresholding parameters for the continuous latent scale are inferred from the data. We used the ordinalNet package [65] to fit the regularized ordinal regression model.

To estimate the seizure intensity for a mouse at each time point of the open-field assay, we fit a hierarchical version of the cumulative link models, namely, cumulative linear mixed models (CLMM). More specifically, we leveraged the repeated measures for each animal in each time bin by treating them as random effects and treated the features tail jerk, leg splaying, side seizure, wild jumping, freezing, and circling as fixed effects. We used leave-out-one-animal cross validation (LOOCV) to validate our ordinal mixedeffects model. We used the estimated model coefficients for both fixed and random effects to estimate the instantaneous per minute seizure intensity for the left-out animal. We used the ordinal package [66] to fit the cumulative linear mixed model.

### 5.8 Model, code, and data availability

The model, code and data will be made available upon publication of the manuscript.

## 6 Author contributions

G.S., L.H., A.M., M.S. and V.K. designed the experiments and analyzed the data. G.S., L.H., and M.M. carried out statistical modeling analysis. All authors wrote and edited the paper.

## 8 Acknowledgments

We thank Kumar Lab members including Brian Geuther, Jacob Beierle, and Jaycee Choi for helpful advice and comments. We thank Erin Day for assistance in carrying out the experiments. This work was funded by The Jackson Laboratory Directors Innovation Fund, National Institute of Health DA051235, DA048634 (V.K.), National Institute of Aging AG078530 (V.K.).

## 9 Supplementary

**Figure S1:**
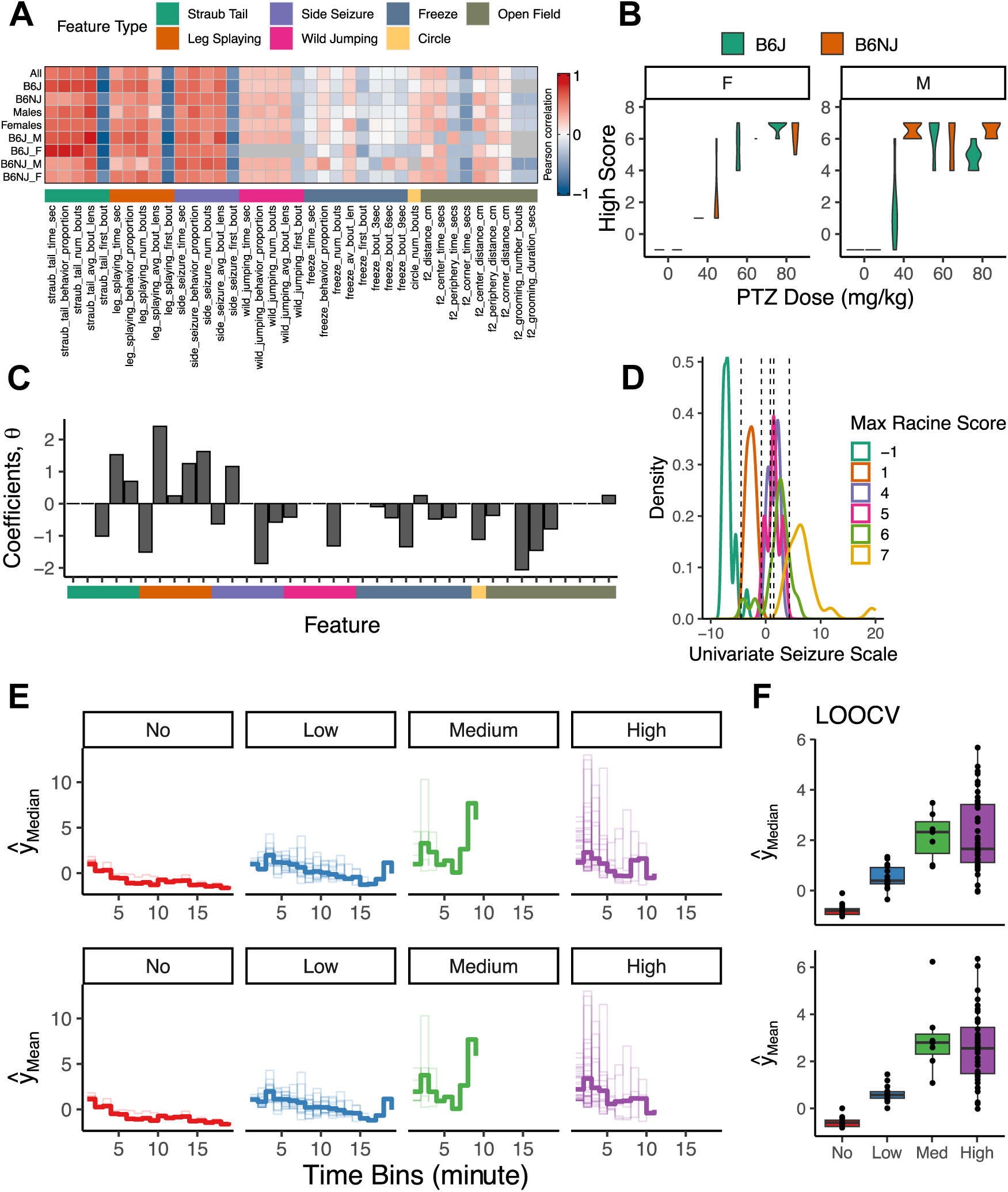
Supplementary data associated with Figures 1, 2, 4. Pearson correlations of features’ summary statistics with dose. (B) A violin plot showing strain differences across females and males. (C) We have plotted the features’ contributions to the seizure intensity scale in (D). (D) We created a univariate seizure scale by training ordinal regression models to create a latent continuous scale with segments corresponding to the original Racine scale scores (Table 1). The resulting distribution of mice along the scale shows an ideal separation of Racine scores except for scores 4, 5, and 6. (E) Instantaneous seizure intensity estimates across 20 1-minute intervals were estimated using supervised hierarchical regression models. The thin lines correspond to individual mice seizure profiles while the bold lines are the median (top) and mean (bottom) respectively, for each seizure group. (F) Validation of the instantaneous seizure intensity model using leave-one-animal cross-validation (LOOCV) where we estimated the median (top) and mean (bottom) seizure intensity of held-out mice (y-axis) and compared it to their true Racine groups (x-axis).

**Table S1:** All scores (single or multiple values) assigned to mice by a scorer. Racine scores of 0 or 2 were not observed. When only the high score was taken into account, no 0, 2, 3 were observed.

| Scores | Highscore | Seizure |
| --- | --- | --- |
| 1 | 1 | Low |
| 4 | 4 | Medium |
| 1, 7 | 7 | High |
| 7 | 7 | High |
| 3, 7 | 7 | High |
| 3, 4-7 | 7 | High |
| 5 | 5 | High |
| 1-6 | 6 | High |
| -1 | -1 | No |
| 4, 6 | 6 | High |
| 3, 7 | 7 | High |

**Table S2:** A table describing the features, their category and units. We calculated open field features, and other behavior features where <BEHAVIOR> = {Straub Tail, Side Seizure, Freeze, Leg Splaying, Wild Jumping, Circle}.

| Video Metrics |  |  |  |
| --- | --- | --- | --- |
| Category | Name | Description | Units |
| Open Field | distance_cm | Sum of locomotor activity | cm |
| Open Field | center_time_secs | Sum of time spent in center | sec |
| Open Field | periphery_time_secs | Sum of time spent along any wall | sec |
| Open Field | corner_time_secs | Sum of time spent in any corner | sec |
| Open Field | center_distance_cm | Average distance from center | cm |
| Open Field | periphery_distance_cm | Average distance from nearest periphery | cm |
| Open Field | corner_distance_cm | Average distance from nearest corners | cm |
| Open Field | grooming_number_bouts | Sum of all grooming bouts | count |
| Open Field | grooming_duration_secs | Average length of grooming bouts | sec |
| Behavior | <behavior>_time_sec | Amount of time the mouse spent in behavior | sec |
| Behavior | <behavior>_behavior<br>_proportion | Proportion of behavior time to total observation<br>time of the experiment | sec |
| Behavior | <behavior>_num_bouts | Number of behavioral bouts for the mouse | count |
| Behavior | <behavior>_avg_bout<br>_lens | Average length of time for a behavioral bout | sec |
| Behavior | <behavior>_first_bout | Time at which the first bout occurs | sec |
| Behavior | freeze_bout_3secs | Number of freezing bouts between 3 and 6 sec-<br>onds | count |
| Behavior | freeze_bout_6secs | Number of freezing bouts between 6 and 9 sec-<br>onds | count |

| Video Features |  |  |  |
| --- | --- | --- | --- |
| Category | Name | Description | Units |
| Behavior | freeze_bout_9secs | Number of freezing bouts greater than 9 seconds | count |
| End of Table |  |  |  |

